# Phylogenomic coupling of F1 chemosensory and archaellum systems across archaea and monoderm bacteria

**DOI:** 10.64898/2026.05.11.724246

**Authors:** Utkarsha Mahanta, Matthew Baker, Gaurav Sharma

**Affiliations:** Department of Biotechnology, Indian Institute of Technology Hyderabad, Sangareddy, Telangana, India; School of Biotechnology and Biomolecular Sciences, University of New South Wales, Sydney, Australia

**Keywords:** Archaellum evolution, chemotaxis, chemosensory systems, phylogenomics, microbial motility, horizontal gene transfer, monoderm bacteria, comparative genomics

## Abstract

Archaellum-associated motility has been viewed as solely archaeal, yet new findings in Chloroflexota prompt a broader perspective. By analysing a curated ∼22,000 NCBI reference genomes alongside 2,397 archaeal and 226 archaellum-encoding Chloroflexota genomes, this study systematically characterises the co-distribution of archaellum loci with chemosensory system (CSS) classes. Maximum-likelihood phylogeny of 3,727 F1-type CheA proteins reveals three major clades, with Clade 1 comprising ∼80% monoderm representation, uniting archaeal and monoderm bacterial lineages in a shared evolutionary grouping. Overall, this work shows that not only archaeal-type motility, but also F1-CSS based sensing system, might have been gained from Archaea to Chloroflexota via horizontal gene transfer and both systems shared an evolutionary trajectory altogether.

## Main

Across prokaryotic evolution, sensing and motility have emerged as mutually reinforcing traits that help organisms cope with fluctuating habitats^1^. To navigate this niche diversity, cells rely on sensory systems that detect chemical gradients, mechanical cues and signals from neighbouring organisms, guiding a variety of motility strategies towards favourable microenvironments^2^. Both processes do not operate in isolation but are coordinated through regulatory pathways that integrate information across multiple scales, allowing cells to adjust their behaviour and motion in real time. Advances in phylogenomics and large-scale genomic studies now reveal how these systems form an interconnected behavioural network that supports rapid, adaptive decisions^3^.

Motility structures such as the flagellum, type IV pili and the archaellum provide the physical means for movement, yet their effectiveness depends on the cell’s ability to detect and interpret environmental cues^4^. Chemosensory systems (CSS) convert external signals into behavioural outputs utilising methyl-accepting chemoreceptor proteins (MCPs), histidine kinases (CheA), and response regulators (CheY)^3^. The coupling between CSS pathway and its cognate motility machinery forms a central decision-making module that allows cells to modulate direction, speed or activation of movement in response to changing conditions. Understanding how these elements are organised and distributed across lineages is therefore essential for interpreting the evolutionary and functional patterns uncovered in comparative genomic analyses^5^.

Although the canonical view holds that bacteria rely on flagella or pili, whereas archaea use the archaellum^6^, recent work by Sivabalasarma et al. indicates that the separation of these systems may not be as strict as once thought^7^. They reported that a few Chloroflexota organisms, including the cultivated species *Litorilinea aerophile*, appear capable of retaining and even using archaellum-related machinery. They also mentioned that only a response regular CheY protein was found in the organism under study, and the CheF protein, which is required for CheY to interact with the archaellum, is completely absent in Chloroflexota and across the bacterial clade.

However, one aspect that must be discussed further is the relative distribution of a similar type of CSS cluster across Chloroflexota and how its presence might have an associated evolutionary link with archaea and their archaellum. Bacteria possess an exceptionally diverse repertoire of chemotaxis pathways, including 17 distinct classes to regulate flagellar motility, one class for type IV pili (TFP) and one class for multiple types of alternative cellular functions (ACF)^8^. In contrast, Archaea predominantly harbour only F1-type chemosensory systems for regulating archaellum-based motility. Monoderm bacterial lineages, i.e., Chloroflexi, and Actinobacteria, along with a few diderm members also encode F1-type chemosensory systems^2,9,10^. The F1 system likely originated from an early lateral gene transfer event from the last bacterial common ancestor during the diversification of Euryarchaeota, enabling newly emerging archaeal groups to expand into novel ecological niches^9^.

To explore these aspects further, we examined 22,204 NCBI-curated reference bacterial and archaeal genomes (Link: https://ncbiinsights.ncbi.nlm.nih.gov/2025/09/02/bacterial-and-archaeal-reference-genome-collection/, downloaded on Nov 5, 2025; 6,850 complete genomes), and used MacSyFinder v2^11^ to identify the genes involved in archaellum, flagellum, type IV pili and Tad system. Sivabalasarma et al. reported 243 draft Chloroflexeota genomes to have archaellum machinery. We additionally included a dataset of 2,397 archaeal genomes to capture all archaeal genomes available in NCBI as of March 23, 2026, of which 369 genomes are complete. Using all three datasets, we further investigated their chemosensory systems by tracking the marker gene CheA, allowing us to evaluate both global and Chloroflexota-specific patterns (Fig. 1, Supplementary Fig. 1 and Supplementary Table 1). It must be noted that all these datasets include draft genome assemblies, along with complete assemblies, owing to which several patterns will stay unidentified unless those genomes are sequenced to completion.

**Figure 1:**
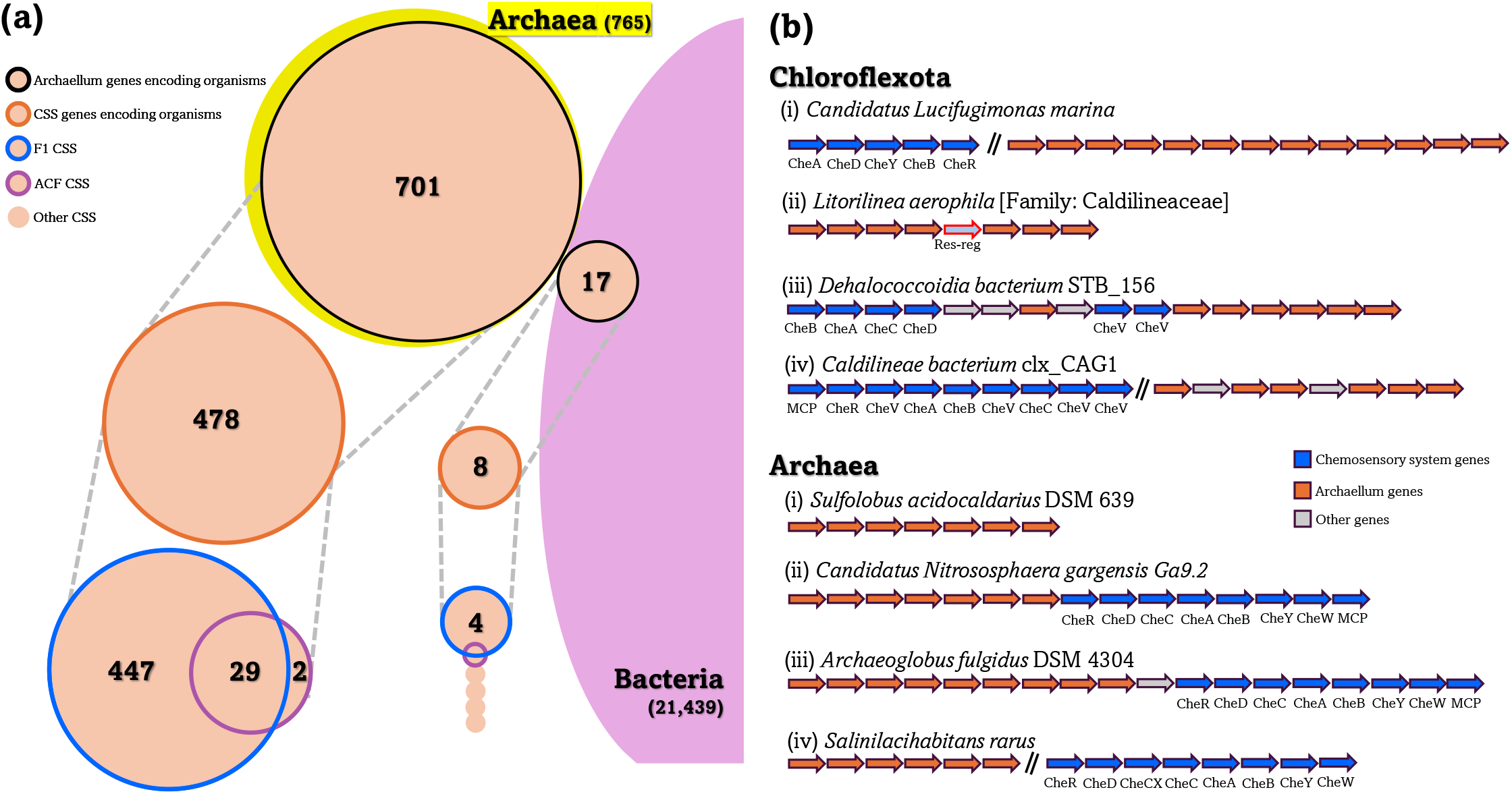
Prevalence and genomic organisation of archaellum and F1-type chemosensory systems, highlighting patterns of co-occurrence, local gene neighbourhoods and synteny between sensory and motility loci. **(a)** Comparative distribution of organisms harbouring archaellum genes and associated F1- and ACF-type chemosensory systems (based on CheA protein) across ∼22k archaeal and bacterial genomes. More than 90% of archaeal genomes in this dataset encode identifiable archaellum components, and the F1-type chemosensory system is the most prevalent sensory module among archaellum-encoding archaea and bacteria. Coloured outlines connected by dotted lines indicate subsets of organisms: black denotes genomes with archaellum genes, orange indicates genomes encoding any chemosensory system, blue marks those with an F1-type system and purple marks those with an ACF-type system. This comparative overview illustrates the uneven yet informative distribution of sensory and motility repertoires across major prokaryotic lineages. **(b)** Representative genomic architectures illustrating the organisation of archaellum gene clusters and the positioning of nearby chemosensory pathways in 22k NCBI and 226 Chloroflexota datsets, highlighting the diversity of locus arrangements across Chloroflexota and Archaea lineages. Within Chloroflexota, panels (i) and (ii) show representatives drawn from the ∼22,000-genome dataset, with panel (ii) corresponding to the organism of interest discussed in the previous study^7^, whereas panels (iii) and (iv) present additional examples from the 226-archaellum-harbouring Chloroflexota subset for which more details have been given in Supplementary table 1C and 1D.

701 out of 765 archaeal genomes contained at least three archaellum genes, consistent with the widespread reliance of archaea on this motility system (Fig. 1a, Supplementary Table 1A). This high prevalence aligns with the classical view that the archaellum is the core and only motility apparatus for many archaeal lineages^2^. The presence of multiple core genes in a genomic synteny strengthens confidence that the detected loci correspond to a functioning archaellum rather than to spurious, isolated homologues. Chemosensory classification based on CheA homologs implied that 476 out of 701 archaellum containing archaea have F1 CSS while 31 organisms harbour an ACF-type CSS, broadly consistent with earlier reports^2^ (Fig. 1a). The comparative enrichment of the F1 class among archaellum-having archaea suggests that specific chemosensory architectures, particularly F1, are preferentially coupled to archaeal motility, apparently due to tighter regulatory or structural compatibility (Fig. 2). Examination of relative genomic synteny showed cases where F1-type CSS loci were immediately adjacent to (eg., *Archaeoglobus fulgidus* DSM 4304, *Archaeoglobus fulgidus*) or located far (eg., *Salinilacihabitans rarus*) from the archaellum genes, indicating lineage-specific variability in sensory and motility genomic organization (Fig. 1b).

**Figure 2:**
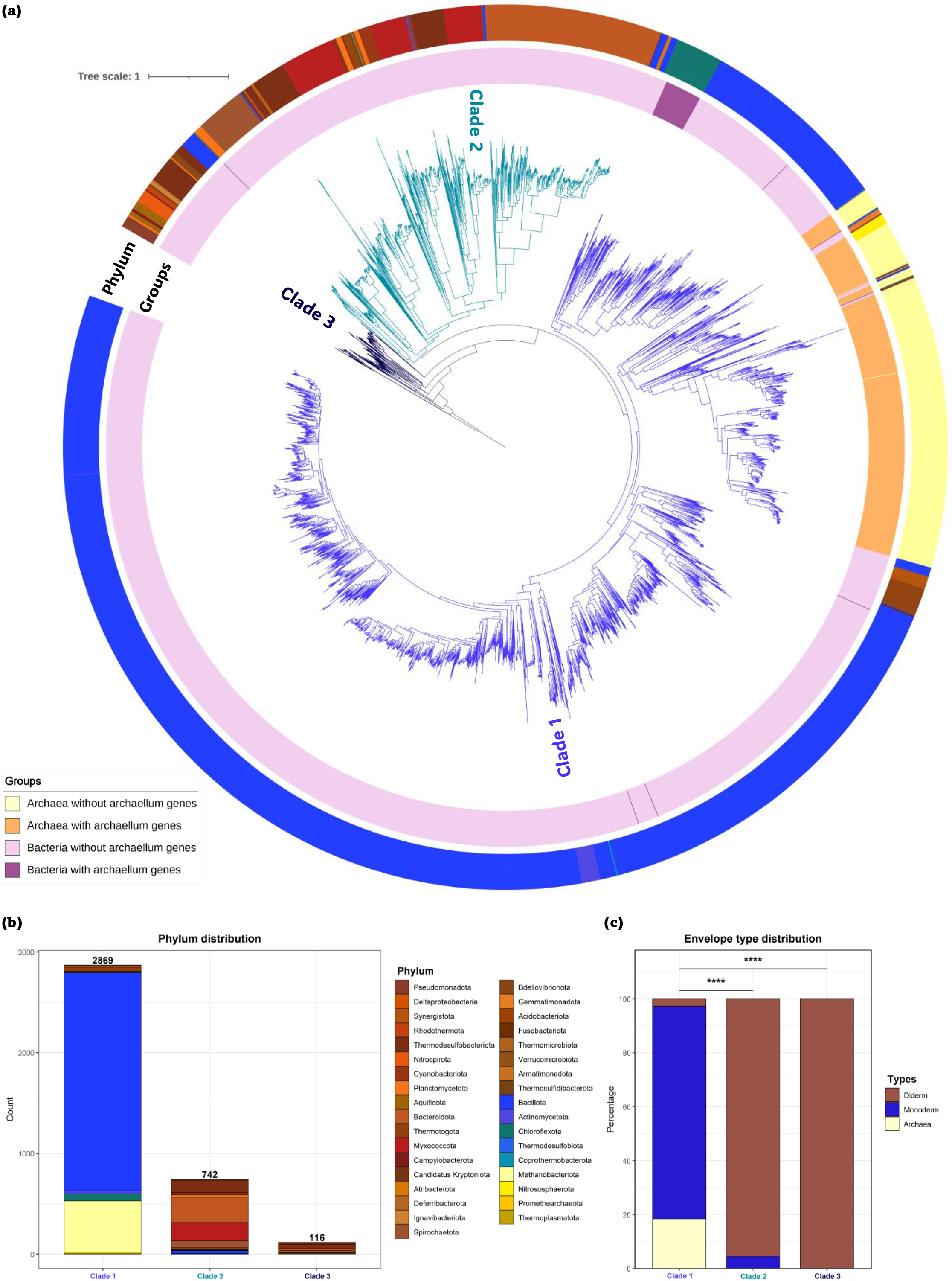
F1-type CheA phylogeny across prokaryotes reveals associations with cell-envelope architecture. **(a)** Maximum-likelihood phylogeny of all 3,727 F1-type CheA proteins identified in the total dataset, reconstructed using FastTree v2. Sequences were aligned with MAFFT and trimmed with TrimAl using the gappyout option, yielding a final alignment of 394 positions. The tree was rooted using an F6-type CheA sequence as an outgroup. The inner coloured ring denotes archaeal and bacterial groups, further distinguished by the presence or absence of archaellum genes. The outer ring indicates phylum-level taxonomy. Three major clades are colour-coded within the F1-type CheA phylogeny. **(b)** Stacked bar plot showing the phylum-level composition of each of the three major F1-type CheA clades, highlighting differences in lineage representation across the tree. **(c)** Distribution of archaeal, monoderm and diderm lineages across the three clades. Statistical comparisons are included to assess the significant enrichment of monoderm taxa in Clade 1 relative to Clades 2 and 3. The details of the data and distribution are provided in Supplementary table 1F.

In stark contrast, and as expected, only 17 out of 21,439 bacterial genomes encoded any archaellum-related genes, reaffirming that bacterial acquisition of archaellum components is extremely rare. Among these, only three Chloroflexota organisms, i.e., *Litorilinea aerophile, Candidatus Lucifugimonas marina* and *Aggregatilinea lenta*, carry recognizable archaellum loci, consistent with the findings of Sivabalasarma et al. (Supplementary Table 1B). Notably, *Candidatus Lucifugimonas marina*, having a genome size of 3.1Mb, encodes a complete F1 CSS positioned 997,160 bp from the archaellum operon but on the same contig, mirroring a similar CSS modular organisation as that within archaea. On the other hand, *Aggregatilinea lenta* carries an ACF-type CSS, likewise co-localised on the same contig and positioned downstream of the archaellum region.

The organism, well-explored by Sivabalasarma et al., i.e., *Litorilinea aerophile*, contains no detectable CSS despite possessing archaellum components. Interestingly, it has a 6.043 Mb draft genome across 93 scaffolds at 98% completeness, raising the possibility that its CSS-genes might be on un-sequenced scaffolds, which must be either experimentally confirmed, or its full genome must be assembled properly. Inspection of all available *Litorilinea* genomes (3–6.5 Mb; all draft-quality) similarly revealed no identifiable CSS components, suggesting that either (a) this lineage genuinely lacks a CSS, or (b) current draft-quality assemblies are insufficiently resolved to capture these loci. Sivabalasarma et al. reported a single response regulator (FKZ61_RS14870) located between archaellum genes in *L. aerophila* and proposed that it may function as a CheY analogue. However, we do not consider this 129-aa protein a plausible CheY candidate: response regulators and CheY proteins are indistinguishable at the domain level, and classification as “CheY” is appropriate only when a regulator is clearly embedded within a CSS. Moreover, MISTdb annotations for this genome report 51 additional response regulators and seven hybrid response regulators, many of which share comparable architecture^12^. Without clear syntenic, functional, or phylogenetic evidence linking FKZ61_RS14870 to a CSS, its designation as a CheY cannot be established.

Beyond the Chloroflexota phylum reported previously^7,13,14^, we also detected archaellum genes in fourteen members of the phylum Bacillota and Actinomycetota, which are monoderm lineages like Chloroflexota, raising the possibility that certain cell envelope structures might be more permissive for the retention and even occasional use of an archaeal type of motility system^15^ (Supplementary Table 1A, 1B). Monoderm organisms, with a single membrane and a thick cell wall, could present a structural context in which archaellum machinery can be more readily maintained or co-opted. However, unlike Chloroflexota species, where flagellum and archaellum genes tend to occur in a mutually exclusive manner, the Bacillota and Actinomycetota genomes that encode archaellum genes also harbour flagellar genes (Supplementary Table 1A). We also found that four of these fourteen bacteria (*Heliorestis acidaminivorans, Anaeromicropila herbilytica, Siminovitchia thermophila*, and *Holtiella tumoricola*) have a F1-type CSS similar to the archaea, which is present a bit far away from archaellum genes in the case of complete genome of *Anaeromicropila herbilytica* or on different contigs in the other three genomes.

Sivabalasarma et al. also reported 244 complete archaellum clusters in Chloroflexota genomes across various orders, following which we also identified their CSS diversity. Out of 226 unique genomes (244 dataset has a few duplicates), 58 organisms harbour F1-type CSS and 24 organisms encode ACF-type CSS, further showcasing the major representation of F1-type CSS in all these archaellum-harbouring Chloroflexota organisms (Supplementary Table 1C, 1D). ACF-type CSS is present significantly in order Aggregatilineales organisms, however, F1-type CSS showed quite sporadic distribution across all orders, including the order Caldilineales, which also has *Litorilinea aerophile*, the organism on which Sivabalasarma et al. has worked. This member, Caldilineae bacterium clx_CAG1 CAG1c_361 (JAAEJZ01; 796 contigs), has both archaellum and CSS clusters on two different contigs, i.e., 361 and 442, further supporting our hypothesis that other order Caldilineales members including *Litorilinea aerophile*, might have a F1-type CSS which should be explored either based on complete genome sequencing or CSS gene amplification. If these systems are not identified based on this, it will imply that this system might have undergone gene loss and there is some other mechanism for archaellum regulation.

Further, statistical analysis supports a significant co-occurrence of the F1 class alongside the archaellum across both archaea and bacteria, specifically monoderm phyla such as Chloroflexota, Bacillota and Actinomycetota, as evident by the Fisher’s exact test: p-value < 2.2e^-16^. This pervasive association strongly supports the hypothesis that the F1 system constitutes the predominant and potentially ancestral sensory architecture governing archaellum-mediated motility across these lineages. While exceptions exist, the strength and consistency of this signal argue against incidental co-distribution and instead point toward functional coupling and co-evolution of these motility and chemosensory modules.

Extending this analysis to core components, we examined the co-occurrence of two conserved archaellum motility genes, *arlI* and *arlJ*, together with the F1-type chemosensory marker gene *cheA* across all three datasets. These three genes were found to co-occur predominantly in members of the archaeal phylum Methanobacteriota, with limited representation in Nitrososphaerota, Thermoplasmatota and Promethearchaeota (Supplementary Fig. 2 and Supplementary Table 1E). Methanobacteriota also represents the most abundant archaeal phylum in both the reference and archaeal datasets (Supplementary Table 1), which may in part contribute to this pattern. In the reference dataset, all three genes were detected in 475 archaeal species and five bacterial species, comprising four Bacillota and one Chloroflexota. Within the Chloroflexota-specific dataset, 57 species encoded all three genes, whereas 726 species in the archaeal dataset carried this complete set (Supplementary Table 1E). It is also important to note that the absence of one or more of these genes in certain genomes may reflect incomplete or fragmented assemblies rather than true biological absence. Phylogenetic reconstruction based on concatenated sequences of these core genes shows that, although Chloroflexota and Bacillota form distinct clades, their sister clades are predominantly composed of Methanobacteriota (Supplementary Fig. 2), consistent with previous observations by Sivabalasarma et al. This pattern further supports a shared evolutionary history and suggests that key components of archaellum-associated motility and its sensory control may have been exchanged between these lineages.

Following this pattern, in-depth phylogenetic analysis of F1-type CheA homologs amidst the whole Bacteria and Archaea dataset further reveals three major clades (Fig. 2a); Clade 1 having archaea grouping together with predominantly monoderm bacterial lineages, whereas Clades 2 and 3 are constituted largely of diderm phyla (Fig. 2 and Supplementary Table 1F). Clade 1 contains 2,869 members, of which nearly 80% are monoderms, a marked enrichment relative to the other clades (Fig. 2b and 2c). The concentration of F1-type CheA in archaea and monoderm bacteria, despite the proposed derivation of monoderms from diderm ancestors^16^, is consistent with a history of horizontal gene exchange between these single-membrane lineages. The repeated association of F1 type chemosensory class with archaellum loci indicates the inheritance of linked modules through gene transfer events. The observed patterns are robust enough to support a more nuanced view of motility and sensing evolution in prokaryotes. How significant their mutual presence is will require transcriptomics and functional studies under different environmental conditions to reveal conditional expression patterns. Further, ecological studies that link motility repertoires to habitat features will clarify how selection shapes the evolution of sensing and movement.

In conclusion, this analysis strengthens the view that sensing and motility are tightly interconnected elements of prokaryotic life, further emphasizing the capacity for evolutionary repurposing and transfer of molecular systems from one kingdom to another. The widespread presence of archaellum genes in archaea and the rare but reproducible occurrences in bacteria reveal a complex evolutionary landscape. The predominant occurence of F1 chemosensory systems with archaellum loci, together with the clustering of archaeal and monoderm bacterial F1-type CheA proteins within a shared phylogenetic clade, hints at a conserved mode of sensory control for this type of motility. Together, these findings invite a future experimental focus on the mechanistic and ecological consequences of coupling between chemosensing and motility machinery, with the promise of uncovering previously hidden strategies that prokaryotes use to navigate their ever-changing world.

## Supplementary data legends

**Supplementary Figure 1: Distribution of archaellum genes and F1-type chemosensory systems across prokaryotes, illustrating their occurrence across bacterial and archaeal lineages**. Maximum likelihood phylogeny was constructed using FastTree v2 from a concatenated alignment of two conserved marker proteins (COG0051: RpsJ and COG0186: RpsQ), comprising 6,308 positions, for the 22k NCBI curated reference bacterial and archaeal genomes. The tree is rooted using the archaeal clade. From the inner to the outer rings, the annotation layers show: phylum-level taxonomy; the number of archaellum genes detected in each genome, shown in red; the number of flagellar genes detected in each genome, shown in green; and the count of F1-type chemosensory components (CheA), shown in blue. All organism details have been provided in the Supplementary Table 1A. Major phylum names have been provided for key representative phyla and organism names have been provided for a few discussed organisms for better understanding.

**Supplementary Figure 2: Maximum-likelihood phylogeny of core archaellum and chemosensory proteins**. The tree was reconstructed using FastTree v2 from a concatenated alignment of ArlI (arCOG01817), ArlJ (arCOG01809) and F1-type CheA sequences. Individual protein sequences were aligned using MAFFT, and the concatenated alignment was trimmed with TrimAl using the gappyout parameter, resulting in a final alignment of 550 positions. The inner ring indicates phylum-level taxonomy, while the outer ring represents class-level classification of the organisms.

**Supplementary Table 1:**

**1A**. Distribution of motility and chemosensory systems across 22,000 bacterial and archaeal genomes. Genes encoding archaellum, bacterial flagellum, T4aP and Tad systems were identified using MacSyFinder v2, while CheA homologues and associated chemosensory signalling modules were detected using hmmscan of HMMER suite 3.4 against the Pfam database v38 along with curated CSS class profiles. Archaeal and bacterial genomes are shaded yellow and purple respectively, to aid visual separation.

**1B**. Chemosensory system profiles stratified by the presence of archaellum genes across ∼22,000 prokaryotic genomes. The analysis distinguishes F1-type and ACF-type signalling modules and compares their occurrence in organisms that encode archaellum components versus those that do not.

**1C**. Distribution of F1-type and ACF-type chemosensory modules among 226 distinct Chloroflexota genomes that contain archaellum genes. These genomes correspond to 244 organisms described in Sivabalasarma et al.; redundant entries were removed to generate a unique set for comparative analysis.

**1D**. Representative examples from the set of 226 Chloroflexota archaellum-encoding genomes, illustrating the genomic neighbourhoods surrounding archaellum loci and their proximity to F1-type chemosensory systems. These cases highlight the possibility of F1 CSS coordinating and interacting with archaellum machinery.

**1E**. The archaeal dataset is presented with the detection status of the three core genes, arlI, arlJ and F1-type cheA. Organisms from the reference and Chloroflexota datasets that encode all three genes are also included here.

**1F**. Comprehensive listing of all F1-type CheA proteins recovered from the dataset and assigned to the three major clades identified in the maximum-likelihood phylogeny. For each entry, the table provides the phylogenetic clade designation, genome accession with locus tag and annotated protein description, phylum-level taxonomy, and broad cell-envelope classification as Archaea, monoderm or diderm.

## Ethics approval and consent to participate

Not applicable

## Availability of data and materials

Data is provided within the manuscript or supplementary information files. Authors have used open-source datasets and tools in this analysis. All tool versions have been provided in the methodology.

## Competing interests

The authors declare no competing interests.

## Funding Declaration

GS acknowledges the Start-up Research Grant from Anusandhan National Research Foundation (ANRF) and IIT Hyderabad for supporting his research. UM is supported by the DST-INSPIRE PhD fellowship, funded by the Department of Science & Technology, Government of India.

## Author Contribution declaration

GS generated the idea. UM performed the analysis and wrote the first draft of the manuscript. MBAB and GS finalized the methodology and validated the results. UM, MABB, and GS edited and finalized the manuscript.

## Consent to Publish declaration

Yes

## Acknowledgement

The support and computational resources provided by PARAM Seva Facility under the National Supercomputing Mission (NSM), Government of India at the Indian Institute of Technology Hyderabad are gratefully acknowledged.

## Notes

### Competing Interest Statement

The authors have declared no competing interest.

## References

1. Beeby, M., Ferreira, J. L., Tripp, P., Albers, S. V. & Mitchell, D. R. Propulsive nanomachines: the convergent evolution of archaella, flagella and cilia. FEMS Microbiol. Rev. 44, 253–304 (2020).

2. Wuichet, K. & Zhulin, I. B. Origins and Diversification of a Complex Signal Transduction System in Prokaryotes. Sci. Signal. 3, (2010).

3. Martín-Mora, D. et al. Functional Annotation of Bacterial Signal Transduction Systems: Progress and Challenges. International Journal of Molecular Sciences 2018, Vol. 19, 19, (2018).

4. Quax, T. E. F., Albers, S. V. & Pfeiffer, F. Taxis in archaea. Emerg. Top. Life Sci. 2, 535 (2018).

5. Gumerov, V. M., Andrianova, E. P. & Zhulin, I. B. Diversity of bacterial chemosensory systems. Curr. Opin. Microbiol. 61, 42–50 (2021).

6. Miyata, M. et al. Tree of motility – A proposed history of motility systems in the tree of life. Genes to Cells 25, 6–21 (2020).

7. Sivabalasarma, S. et al. Structure of a functional archaellum in Bacteria of the Chloroflexota phylum. Nature Microbiology 2025 10:10 10, 2412–2424 (2025).

8. Ortega, D. R. et al. Repurposing a chemosensory macromolecular machine. Nature Communications 2020 11:1 11, 2041- (2020).

9. Briegel, A. et al. Structural conservation of chemotaxis machinery across Archaea and Bacteria. Environ. Microbiol. Rep. 7, 414–419 (2015).

10. Ortega, D. & Beeby, M. How Did the Archaellum Get Its Rotation? Front. Microbiol. 12, 803720 (2022).

11. Néron, B. et al. MacSyFinder v2: Improved modelling and search engine to identify molecular systems in genomes. Peer Community Journal 3, (2023).

12. Gumerov, V. M., Ulrich, L. E. & Zhulin, I. B. MiST 4.0: a new release of the microbial signal transduction database, now with a metagenomic component. Nucleic Acids Res. 52, D647–D653 (2024).

13. Hug, L. A. et al. Community genomic analyses constrain the distribution of metabolic traits across the Chloroflexi phylum and indicate roles in sediment carbon cycling. Microbiome 2013 1:1 1, 22- (2013).

14. Lim, Y., Seo, J. H., Giovannoni, S. J., Kang, I. & Cho, J. C. Cultivation of marine bacteria of the SAR202 clade. Nature Communications 2023 14:1 14, 5098- (2023).

15. van Wolferen, M., Pulschen, A. A., Baum, B., Gribaldo, S. & Albers, S. V. The cell biology of archaea. Nature Microbiology 2022 7:11 7, 1744–1755 (2022).

16. Witwinowski, J. et al. An ancient divide in outer membrane tethering systems in bacteria suggests a mechanism for the diderm-to-monoderm transition. Nature Microbiology 2022 7:3 7, 411–422 (2022).

