## Supplementary figures and images for "Phylogenomic coupling of F1 chemosensory and archaellum systems across archaea and monoderm bacteria"

### Supplementary Fig. 1

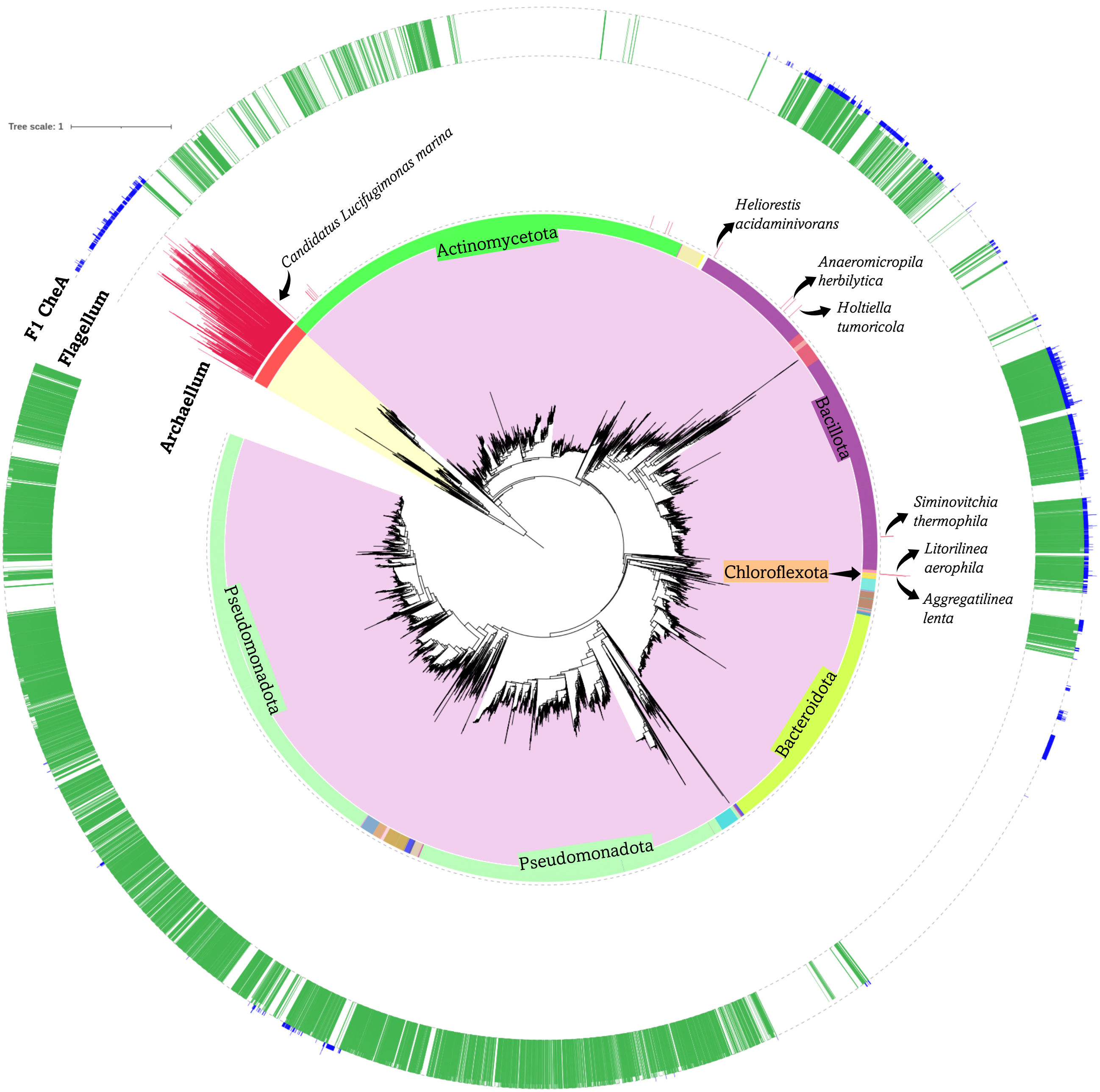

### Supplementary Fig. 2

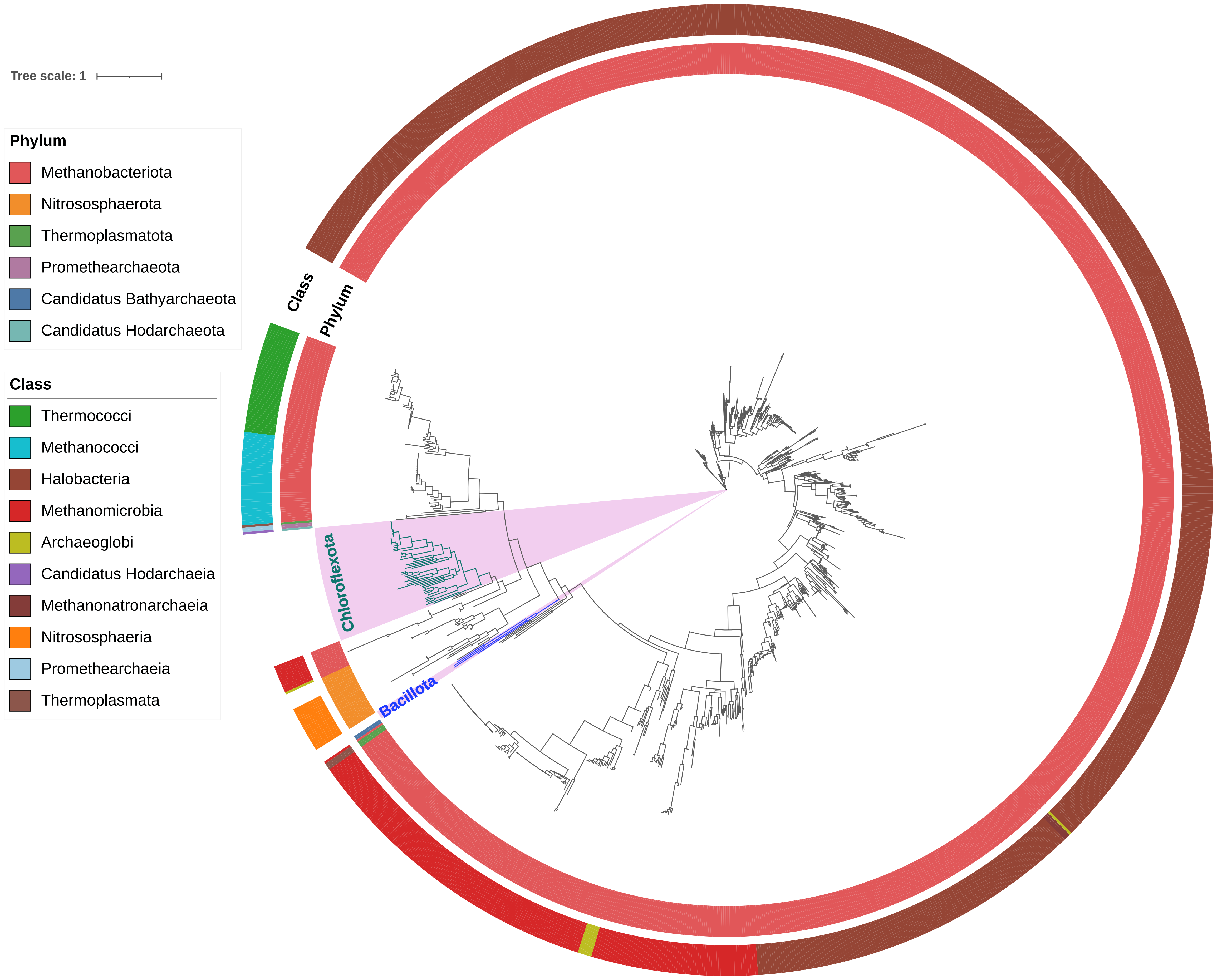
